# The plasma miR-122 basal levels respond to circulating catecholamines in rats

**DOI:** 10.1101/257402

**Authors:** Xu Peng, Qiao Li, Song Lu, Xueling He, Sisi Yu, Zhihui Zhang, Guohui Xu, Lu Li, Tinghan Yang, Jiang Zhu, Wenli Zhu, Zhigang Wu, Delun Luo, Jie Zhu, Binghe Xu, Jian Huang, Hailin Yin, Kai Xu

**Affiliations:** The Laboratory Animal Center, Sichuan University, Chengdu, Sichuan, 610041, China; National Cancer Institute/Department of Medical Oncology, Cancer Hospital, Chinese Academy of Medical Sciences and Peking Union Medical College, Beijing, 100021, China; Sichuan Cancer Hospital & Institute, Sichuan Cancer Center, School of Medicine, University of Electronic Science and Technology of China, Chengdu, Sichuan 610041 China; West China Hospital, Sichuan University, Chengdu, Sichuan 610041 China; TCM Symptons and Precisional Medicine, Chengdu University of Traditional Chinese Medicine, Chengdu, Sichuan, 610075, China; Center for Informational Biology, University of Electronic Science and Technology of China, Chengdu, Sichuan 610041 China.

**Keywords:** hsa-miR-122-5p, catecholamine, drug-induced liver injury, biomarkers

## Abstract

miR-122 in circulation is a promising non-invasive biomarker as a replacement or supplement of current serum biomarkers for liver injuries. But the concept was questioned by recent studies, mainly due to its release from hepatocytes in absence of overt cellular injuries. In this study, we reported that the hepatic metabolism of circulating catecholamines resulted in the release of hepatocyte-specific miR-122. Acute stress-induced hepatocellular deformation was histopathologically different from drug-induced liver injury with significant increases of plasma miR-122 levels. The basal levels of human plasma miR-122 could be significantly altered by emotional responses. Interday variances of plasma miR-122 measurements were reduced effectively by stress-relief measures. The metabolism of basal circulating norepinephrine and epinephrine in liver might contribute to the basal levels of plasma miRNAs expressed in hepatocytes.

## INTRODUCTION

Drug-induced liver injury (DILI) accounted for large portions of clinic mortalities due to acute liver failure and idiosyncratic DILI is the common cause for unfavorable drug withdrawals in western countries(Shi, *et al* 2013). In Asian, herbal and dietary supplements were the most common causes of DILI(Wai, *et al* 2007). Serum alanine aminotransferase (ALT) is the current standard biomarker for DILI, but it suffered in liver-specificity, narrow detection range and long circulating half-life as reviewed(Kim, *et al* 2008). MicroRNA (miRNA) is a family of short noncoding transcripts (22 nucleotides in length) involved in negative post-transcriptional regulation of mRNA through translational repression and cleavage(Bartel 2009, Lee, *et al* 1993, Xu, *et al* 2016). A valuable collection of hundreds of protein- or exosome-associated miRNAs is shed into blood from organs or tissues under physiological or pathological events(Chen, *et al* 2008, Mitchell, *et al* 2008, Wang, *et al* 2009). The miRNA biogenesis is cell-associated and event-driven that makes the miRNA expression profile an indicator of cell types or signal pathways(Bartel 2009). Those features brand plasma miRNAs the ideal non-invasive biomarkers for pathological changes. Plasma miRNAs originated from multiple types of cells and released under various events, which complicated the tasks to define their basal levels(Witwer and Halushka 2016). Despite the extensive studies on plasma miRNAs, their release mechanisms, basal levels, metabolic rate and other pre-requisite parameters for measurement are still accumulating.

The expression of miR-122 exhibits a hepatocyte-specific pattern in vertebrates, with content up to 70% of all hepatocellular miRNAs(Chang, *et al* 2004, Ludwig, *et al* 2016). Changes of miR- 122 and miR-192 concentrations in circulation demonstrated strong correlations with plasma ALT and liver histology in mouse models in acetaminophen (APAP)-induced liver injury (AILI)(Wang, *et al* 2009). Plasma miR-122 elevations were detected significantly earlier than plasma ALT changes, and plasma miR-122 was suggested as a potential biomarker for the early onset of DILI. The predictive value of plasma miR-122 for liver injury was later confirmed in rats(Sharapova, *et al* 2016, Starckx, *et al* 2013, Yamaura, *et al* 2012), monkeys and human subjects(Cermelli, *et al* 2013, Starkey Lewis, *et al* 2011, Starkey Lewis, *et al* 2012). MiRNA profiling in DILI and ischemic patients revealed that a set of 11 circulating miRNAs (including miR-122) that could discriminate AILI from ischemic hepatitis(Ward, *et al* 2014). Differential expressions of plasma miR-122 were observed and confirmed in patients with chronic liver diseases, such as viral-related hepatitis(Cermelli, *et al* 2011, Waidmann, *et al* 2012, Yamaura, *et al* 2017, Zhang, *et al* 2010), cholestasis, steatosis(Yamaura, *et al* 2012) and fibrosis(Halasz, *et al* 2015, Trebicka, *et al* 2013). Attempts were also made to use plasma miR-122 as predictive biomarkers for the onset and clinic outcomes in patients with acute liver diseases(Fontana, *et al* 2014, Wang, *et al* 2012, Ward, *et al* 2014). Compared to the above-mentioned researches, more studies were carried out on plasma miR-122 as non-invasive DILI biomarker(Singhal, *et al* 2014, Starkey Lewis, *et al* 2011, Su, *et al* 2011, Wang, *et al* 2009, Yang, *et al* 2015).

Regardless of detection methods and normalization approaches, plasma miR-122 was consistently detected from all healthy subjects from rodents to human, with extremely large intra- and interindividual variances. In a recent effort to utilize capillary blood for plasma miR-122 measurements aimed at point-of-care DILI test, the plasma miR-122 concentrations of the finger-prick blood drops from different fingers varied in a range of 4 to 6 folds intra-individually in most subjects, and additionally, ~20-fold changes in 2 cases were observed between fingers (Vliegenthart*, et al* 2017). It was difficult to dismiss those differences as technical variances. While efforts were made to reduce the intra-individual variances, studies of plasma miR-122 as DILI biomarkers were reported recently in more negative tones in general, due to its release from hepatocytes in absence of overt cellular injuries(Church and Watkins 2017). Additionally, the diurnal variation of plasma miR-122(Heegaard, *et al* 2016) is also a concern for its intraday variance, especially those during daytime. In an attempt to understand what contribute to the high variation of circulating miR-122 measurement, we utilized plasma-direct miRFLP assays(Xie, *et al* 2015, Zhu, *et al* 2017) and stress-trained rat models to explore the effects of stress and drugs on plasma miR-122 levels.

## RESULTS

### Inconsistency between liver histology and plasma miR-122 levels in AILI mice

To investigate the correlation of liver histopathology and plasma miR-122 concentrations, 68 BALB/c mice were treated with different doses of APAP. The plasma miR-122 concentrations in samples collected at different time points were analyzed *(Figure 1A).* The plasma miR-122 concentration of untreated mice was measured at 9,704 ± 2,406 copy (cp)/μl (Mean ± SEM, n = 4) with a narrow inter-individual coefficient of variation (CV) of 37.9%. The average plasma miR- 122 concentration of saline-injection group, to our surprise, was 155,695 ± 23,450 cp/μl (Mean ± SEM, n = 4) with inter-individual CV of 28.7%, a 16-fold increase over that of untreated group *(p* = 0.008). The plasma miR-122 levels in mice of low-dose groups peaked between 8 to 24h, with large inter-individual CV ranges of 84.4% to 331.4%. Ironically, the average plasma miR-122 levels from mice in medium- and high-dose groups were significantly lower than that of low-dose groups with narrower inter-individual CV ranges of 40.0% to 240.9%. The quantitative analysis revealed the deviations from the dose-dependent correlation between APAP administration and plasma miR-122 levels. The results suggested the involvement of additional confounding factors other than APAP, possibly the acute stresses caused by saline injection and blood collections (needle punctures). By inhibiting cyclooxygenases in central nervous system, APAP increased pain threshold with reduced awareness in treated mice. This could explain the narrower interindividual CV ranges in medium- and high-dose APAP groups at 2, 8 and 24h.

**Figure 1.**
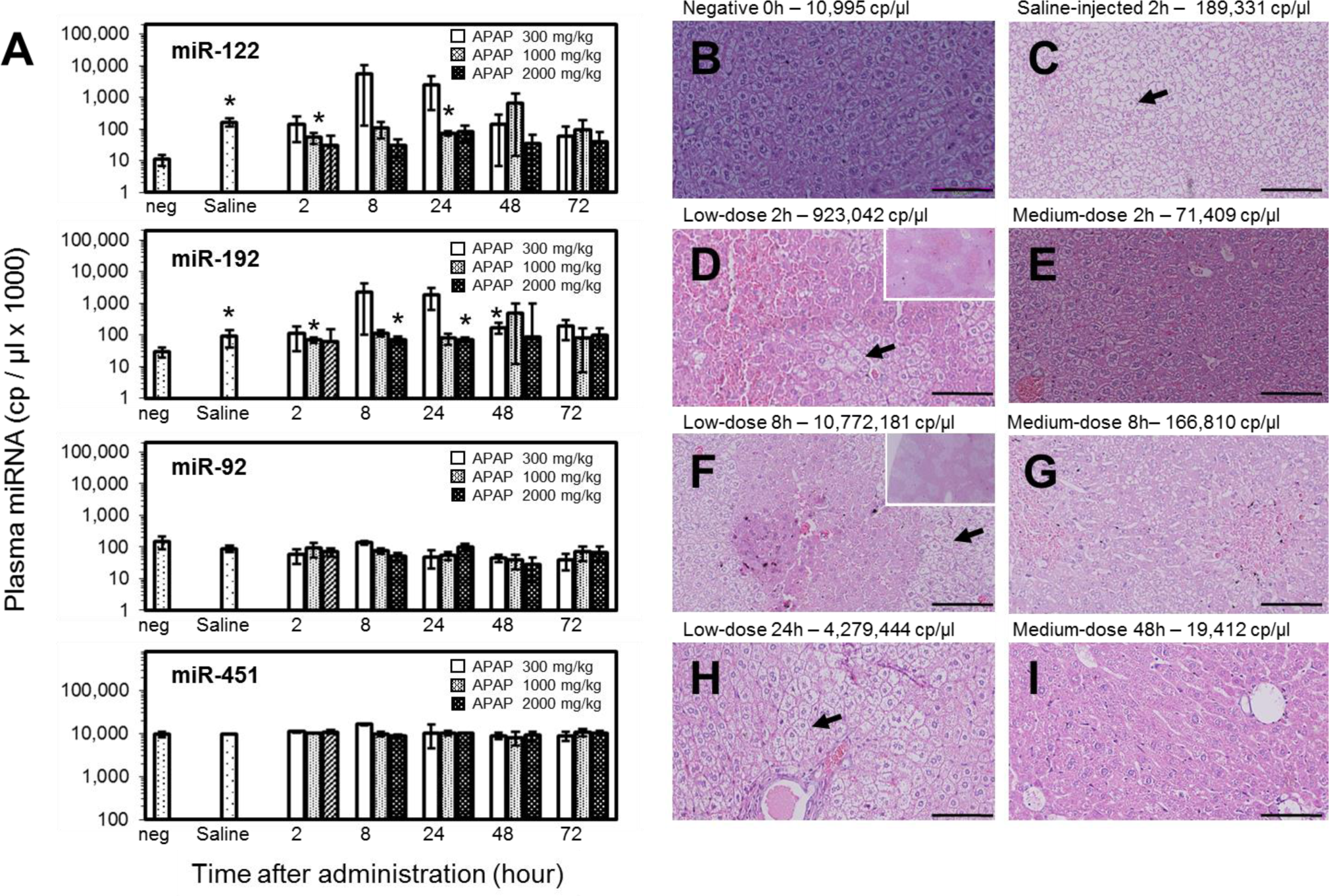
Time-courses of plasma miRNAs and liver histology in mice induced by APAP. (A) The temporal measurements of plasma miRNAs (miR-122, miR-192, miR-92, and miR-451) from mice treated by APAP. (n = 4/group). Stars indicate the measurements with significant difference from negative mice (p < 0.05) (Mean ± SD). The representative histopathological changes are shown for: (B) negative rat with normal histology; (C) rat with saline-injection 2-h before sacrifice; (D) rat at 2h in low-dose APAP group. Spotted hemorrhage lesions were shown in small frame; (E) rat at 2h in medium-dose APAP group; (F) rat at 8h in low-dose APAP group. Spotted hemorrhage lesions were shown in small frame; (G) rat at 8h in medium-dose APAP group; (H) rat at 24h in low-dose APAP group. (I) rat at 48h in medium-dose group. The histopathological features of enlarged hepatocytes and nucleus shifts were indicated by arrows. H&E x 200. Scale bars = 100 μm.

Among the miRNAs measured, miR-192 is also enriched in rodent hepatocytes and its plasma concentrations responded to APAP treatment in correlation with the plasma miR-122 significantly (Pearson’s coefficient: 0.957, *p* < 0.01, 2-tailed). The plasma concentrations of miR-451 (Red blood cell-associated) and miR-92 (ubiquitous expression) were used as internal quality assurance.

### Two distinct types of liver histopathology in low-dose AILI mice

Histopathological analysis were performed on mouse liver sections and the representative results were listed *(**Figure 1 B to I**).* In correlation with the 16-fold increase of plasma miR-122 over the mean value of untreated controls, liver histology of the saline-injection mice presented unique patterns of loose tissue structures, enlarged hepatocytes, and nucleus shifts in the absence of overt inflammatory cell infiltration and necrosis/hemorrhage *(**Figure 1C**).* In low-dose APAP groups, higher plasma miR-122 were observed initially (at 2, 8 and 24h) with larger inter-individual variation. The enlarged hepatocytes and nucleus shifts, similar to that of saline-injection mice, were observed in mice with high plasma miR-122 concentration *(**Figure 1D, F and H**).* In addition, centrilobular expansion and congestion/necrosis were spotted in correlation to the overexpressions of plasma miR-122. In medium-dose APAP groups, plasma miR-122 levels of 20,430 to 71,409 cp/μl were determined for mice at 2h in the absence of overt histopathological changes. In 8h medium-dose group, centrilobular abnormality and loose tissue structures appeared in 2 mice which had plasma miR-122 concentrations of 166,810 and 119,784 cp/μl. The loose tissue structures, necrosis and hemorrhage, were observed for mice in 48h medium-dose group regardless of their plasma miR-122 levels (1,310,081, 2,029,503, 19,412, and 5,764 cp/μl).

Overall, the data suggest that there were two distinct histological types of hepatocellular alterations. One type was represented by necrosis of hepatocytes caused by DILI with sustained elevations of plasma miR-122. Another type was the stress-induced Hepatocellular Deformation (siHD), represented by enlarged hepatocytes, nucleus shifts in absence of noticeable necrosis with instantaneous miR-122 releases in responses to physiological stressors, such as saline injection, blood collection, hemorrhage and animal death etc (Hart, *et al* 1989).

### The dynamic plasma miR-122 levels revealed siHD in Cisplatin-treated rats

Cisplatin (CDDP) is a common chemotherapeutical reagent without known links to stress suppression, so that it was selected to verify that if siHD could be induced in naïve rats by acute stresses caused by frequent blood samplings. Two baseline samples were collected 2 days before stress stimulation and saline or CDDP treatment and frequent blood samplings were performed at 48h time mark. The plasma miR-122 concentrations and representative liver histological sections are illustrated (***Figure* 2**). For the saline control group, the miR-122 levels peaked quickly with great inter-individual variation (44,023 to 247,184 cp/μl), and returned to basal levels within 2 h *(**Figure 2 A to C**).* When the same operations were repeated at 96h, significantly reduced plasma miR-122 levels were observed. It implied that these rats adapted to the stresses of saline injection and frequent blood samplings. In CDDP-treated groups, sharp peaks were observed in 8 rats immediately after the first treatments at 48h regardless CDDP dosages *(**Figure 2 D to K***), and followed by sustained plasma miR-122 elevations which returned slowly to basal levels. After second CDDP administrations at 96h, sustained plasma miR-122 levels were determined in all CDDP-treated rats and 6 out of 9 rats died before the 144h endpoint. The biopsies of liver sections showed necrosis of hepatocytes in CDDP-treated rats *(**Figure 2 E-1, H-1 and J-1**)*, enlarged but not necrotic cells in negative control rat ‘A’ *(**Figure 2A-1**)* and normal liver histology in negative control rats ‘B’ and ‘C’ (data not shown). Parallel comparison of plasma miR-122 levels between saline and CDDP-treated mice, revealed distinct peaks for siHD and flattened slopes for DILI, respectively *(**Figure 2 B and E**)*. siHD and DILI could be distinguished by temporal measurement of plasma miR-122 after stress eroding.

**Figure 2.**
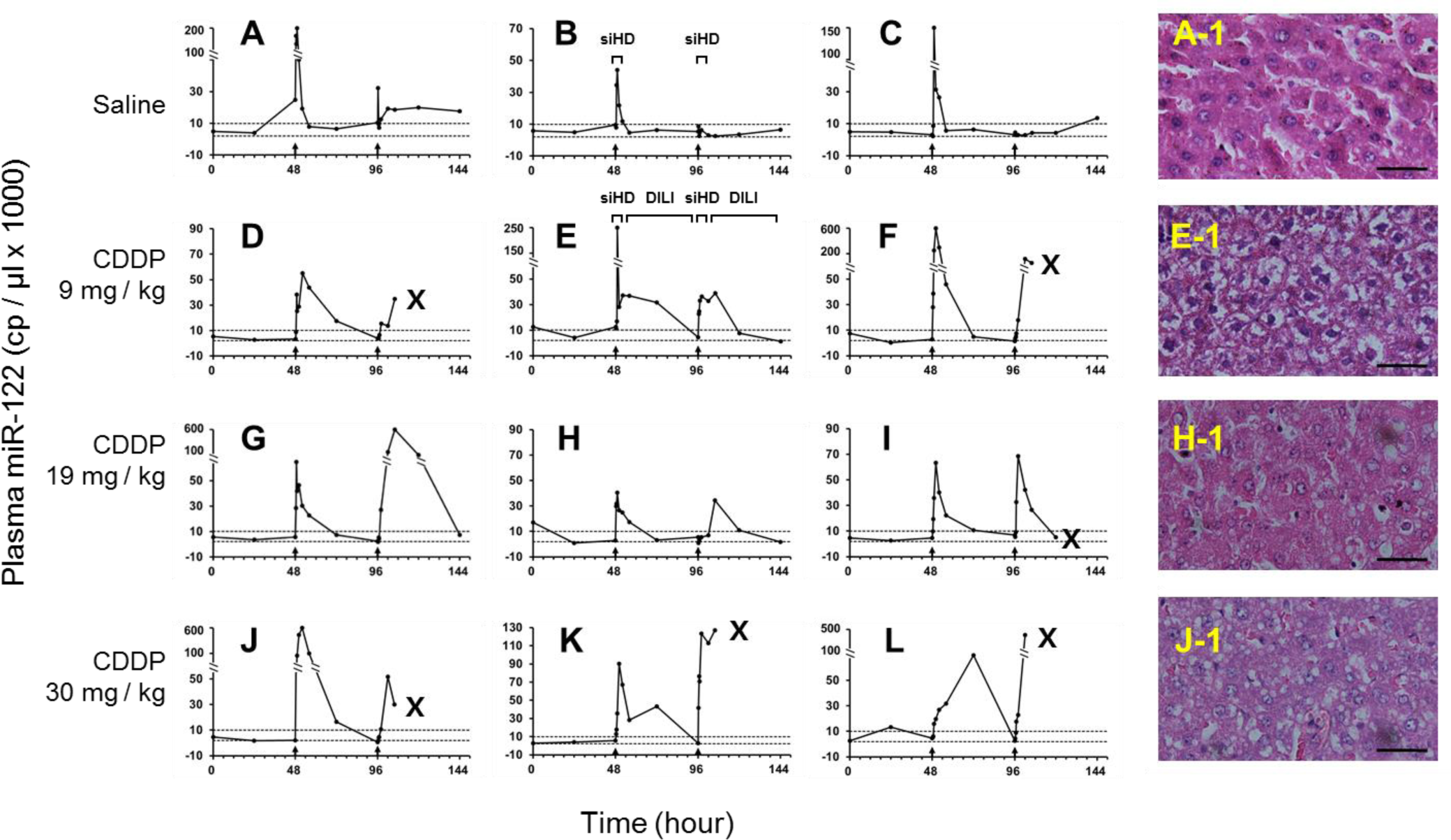
The profiling of rat plasma miR-122 levels after CDDP treatments at 48h and 96h respectively. The miR-122 concentrations from rat plasma samples collected before and after two i.v. administrations of (A-C) saline; (D-F) 0.9 mg/kg; (G-I) 1.9 mg/kg; (J-L) 3.0 mg/kg of CDDP, respectively. The administrations of medications at 48h and 96h are indicated by upper arrows and followed by blood collections before and at 20, 40, 60, 120, 240, and 480 min after treatments. X marks animal death; (A-1) histopathological features of negative control rat ‘A’, showed enlarged hepatocytes and nucleus shifts; (E-1) histopathological features of rat ‘E’, showed necrosis/apoptosis; (H-1) histopathological features of rat ‘H’, showed necrosis/apoptosis and vacuolation; (J-1) histopathological features of rat ‘J’, showed necrosis/apoptosis and vacuolation. H&E x 400. Scale bars = 100 μm.

### Catecholamines promoted plasma miR-122 levels in stress-trained rat models

Six S/D rats were trained with stress-training routines as described in Materials and Methods to overcome the stressful influence by activities of injection and blood collections on plasma miR- 122 measurement. The drugs administered and plasma miR-122 measurements are illustrated *(**Figure 3 A to C**).* The duplicate tests were displayed in Supplementary Materials *(**Figure S1**).* The plasma miR-122 concentrations of the negative control rats #5 and #6 were averaged at 3,163 ± 240 cp/μl and 4,347 ± 279 cp/μl (Means ± SEM, n = 40) from Day 1 to Day 22, with statistical difference *(p* = 0.003, One-way Anova). When grouped by the 4 sampling times during daytime individually, the plasma miR-122 concentrations showed no difference in the same period *(**Figure 4A**).* This demonstrates that the basal levels of plasma miR-122 during daytime were relatively stable for a long period in stress-trained rats during daytime. It implied that the daytime basal levels of plasma miR-122 might be definable individually under stress-free conditions. The same trend was confirmed by statistics from other four rats in the untreated training periods (data not shown). No weight loss or abnormal behavior issues observed for rats #5 and #6 during the 22-day stretch and the frequent blood samplings were well tolerated in the rat models.

**Figure 3.**
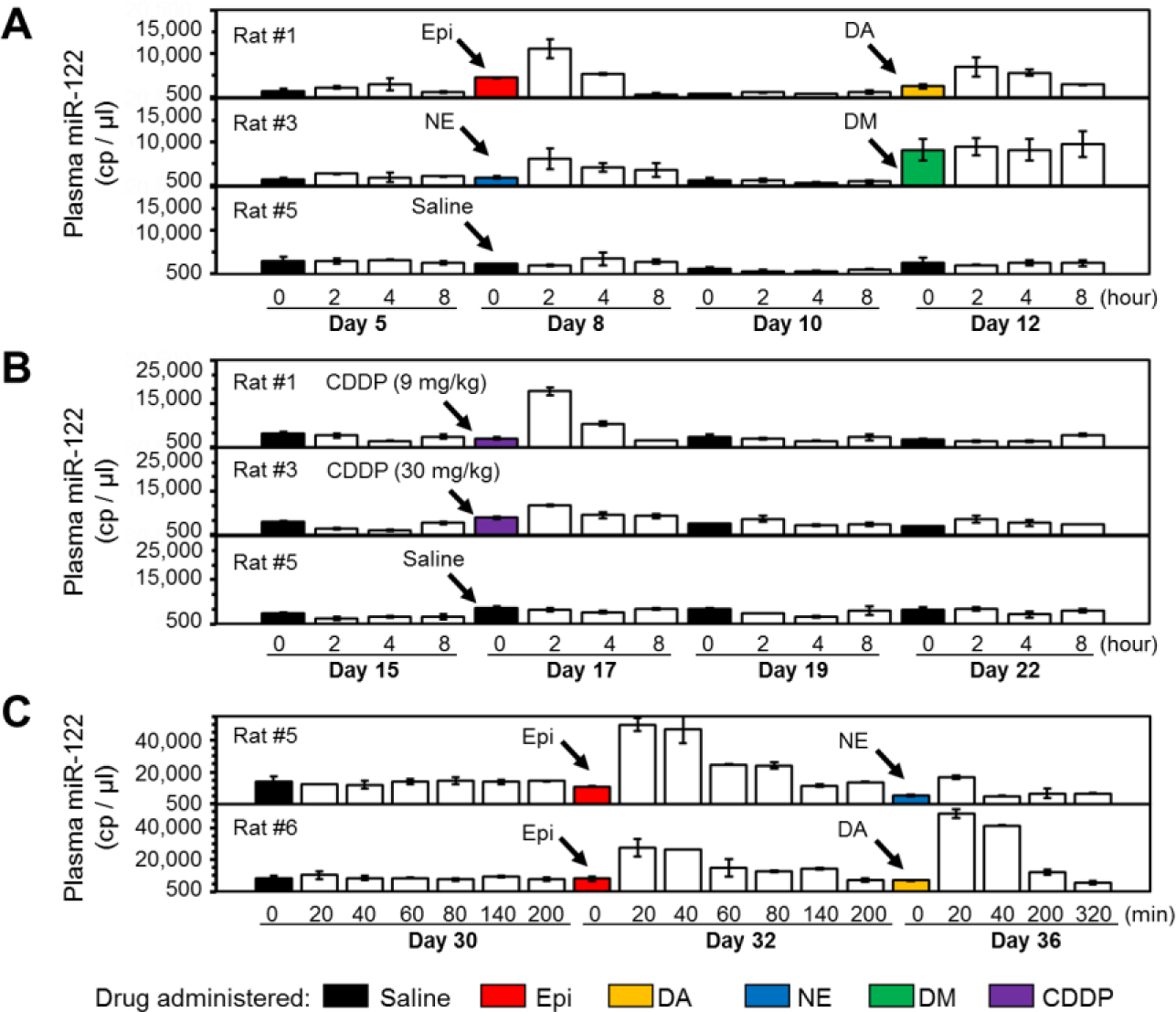
Plasma miR-122 expressions by Catecholamines and CDDP in stress-trained rats. (A) Plasma miR-122 changes before and after the administrations of Epi, DA, NE, and DM; (B) Plasma miR-122 changes before and after the administrations of CDDP at 9 mg/kg and 30 mg/kg respectively; (C) Plasma miR-122 alteration in blood samples collected in shorter sampling intervals. 100 |il of solution containing 0.1 mg of Epi (red box), 0.2 mg of NE (blue box), 1 mg of DA (orange box), 0.5 mg of DM (green box), and saline (black box) were i.v. administrated at indicated time point, respectively. Mean ± SD, n = 3.

After i.v. administrations of Epi, NE, dopamine (DA) or dexamethasone (DM), significant elevations of plasma miR-122 were observed in stress-trained rats (***Figure* 3A**). In concert with short circulating half-lives of all three catecholamines (2 to 3.5 min)(Goldstein, *et al* 2003), the clearance of plasma miR-122 was observed within 2 h after administration. Meanwhile, sustained plasma miR-122 elevations were detected in rat plasma samples 8 h after DM treatment in corresponding to the 190 minute (min) half-life of DM in circulation(McEwen 2008). At shorter sampling intervals, a peak of 3.85-fold increase of plasma miR-122 by Epi administration was observed which returned to basal levels in 200 min. The numbers were 6.62-fold increases in 200 min for DA and 2.97-fold increases in 40 min for NE *(**Figure 3C**).*

In stress-trained rats treated with CDDP, plasma miR-122 peaks between 10,568 to 35,288 cp/μl were obtained at 2h after drug administration, but returned to basal levels at 8h except for one rat (rat #2), which was apparently stimulated emotionally before treatment for unknown reason *(**Figure 3B and Figure S1B**).* Those values were significantly lower than the plasma miR-122 levels at 8h after CDDP administration in naïve rat models (Table S1). It implies the potential synergistic enhancement of CDDP-induced liver injuries by stress sensitization in naïve rats.

### Psychological stress was the predominant contributor to miR-122 release

To ascertain the role of stress in miR-122 release, stress stimulating tests were performed on six naïve female S/D rats as described in Materials and Methods. The most notable contributor that stimulating the significant plasma miR-122 upregulation was the order of treatment received by individual rats *(**Figure S2**).* Even before any treatment, an average 2.69-fold elevation of plasma miR-122 was observed in 3 samples collected subsequently over their counterparts collected first in each group. The largest plasma miR-122 increase of 125.0-fold over basal level was observed in rat with saline injection without ether-inhalation after 4 blood samplings *(**Figure 4B**)*, comparing with the 22.5-fold increase in rat with ether-inhalation. Overall, our data suggest that the primary contributor to hepatocellular miR-122 release was acute stress and the light ether sedation reduced miR-122 release dramatically. Further comparison demonstrated that the stress-training routines reduced plasma miR-122 variances significantly *(**Figure 4C**).*

**Figure 4.**
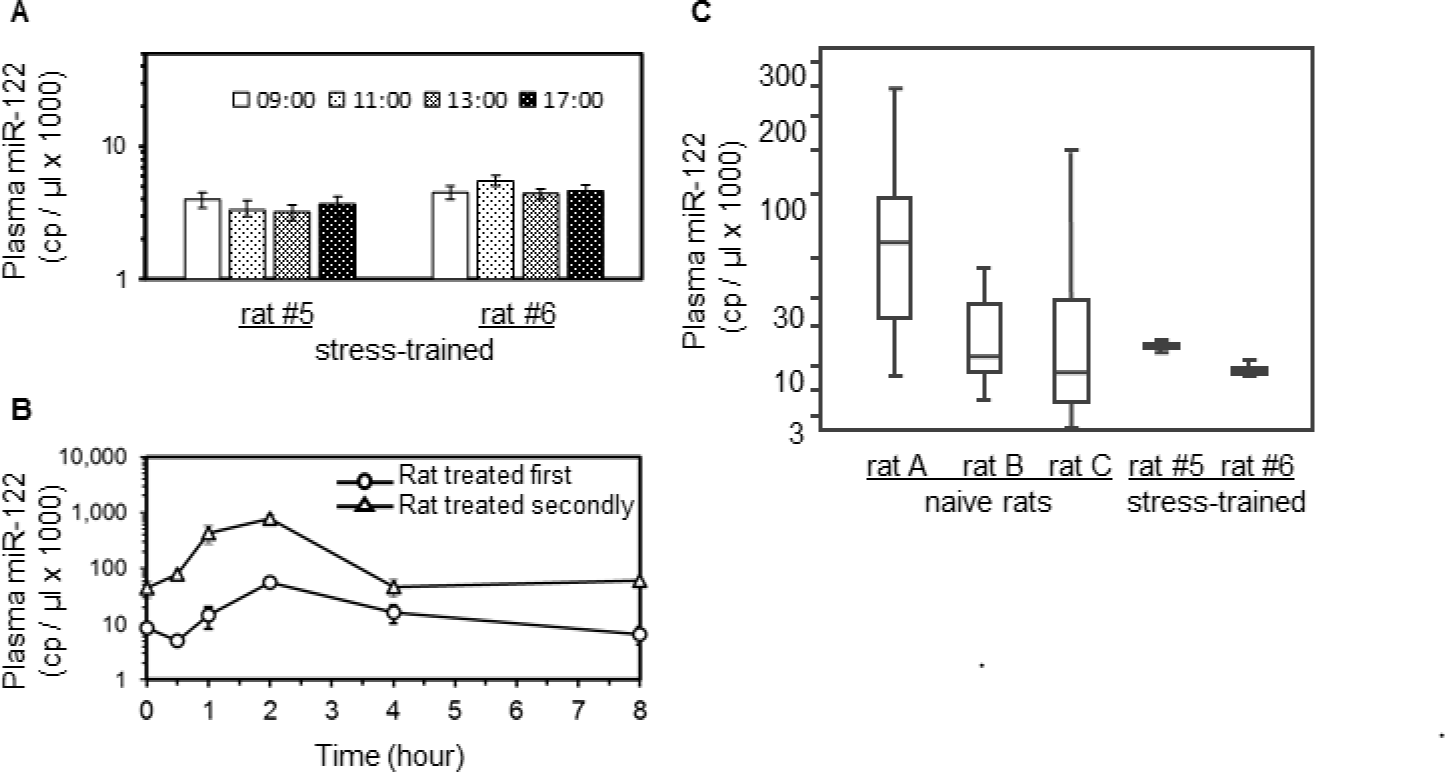
Plasma miR-122 expressions induced by stress. (A) The miR-122 concentrations from plasma samples collected at different times in a given day during a stretch of 22 days (n = 10) on stress-trained rats #5 and #6 (Mean ± SEM); (B) The influence of frequent blood collections on plasma miR-122 concentrations in two naïve rats by repeat blood sampling at 0, 0.5, 1, 2, 4, and 8h; (C) The comparison of 7 plasma samples collected in shorter sampling intervals (at 48h to 52h) from naïve rats ‘A’, ‘B’, and ‘C’ *(**Figure 2 A to C**)*, against those from stress-trained rats #5 and #6 on Day 30 *(**Figure 3C**).* n = 3.

To compare the relationship between plasma concentrations of catecholamines and miR-122, two hundred microliters of blood samples were collected from 4 naïve female S/D rats (7-week-old) at 0.33, 0.66, 1, 2, and 4h, respectively, without ether sedation. Plasma concentrations of miR-122 along with brain-specific miR-9 and miR-124 were determined by plasma-direct miRFLP assay. Plasma concentrations of NE, Epi and DA were determined by ELISA assay from 3 rats with enough plasma samples. The relationship between the plasma contents of miR-122 and three catecholamines were analyzed *(**Figure S3 A to D**)*. Among the four rats, two had plasma miR-122 peaks over 10-folds of their basal levels and another two had constant elevations of plasma miR- 122 (> 50,000 cp/μl). While NE concentrations were maintained constantly at ~500 pg/ml in all tested samples, the plasma concentrations of Epi and DA were fluctuated dramatically. The venous plasma concentrations of miR-122 were negatively correlated with the venous plasma Epi (Pearson’s correlation: *γ* = -0.954, *p* = 0.012 and *γ* = -0.939, *p* = 0.018, respectively) when the measurements at 0h were removed as outliners. But when the first sets of data were included, Pearson’s corrections became insignificant *(γ* = -0.680, *p* = 0.137 and *γ* = -0.777, *p* = 0.069, respectively). This dramatic changes in correlation relationship implied that there were discordance in event sequences between the first sets and the rest of the samples. No significant correlations between miR-122 and Epi were observed in other two rats which had constantly elevated plasma miR-122.

### The metabolic rates of miR-122 in circulation

The metabolic rates of plasma miR-122 induced by different reagents were calculated and summarized in Table 1. The mean metabolic rate of plasma miR-122 was 43.2 ± 15.9 min in rats during the first siHD period, then shrank to 18.1 ± 11.2 min in the second siHD period with minimal of 11.1 and 12.0 min in naïve rat models *(**Figure 2 A to C**).* The metabolic rates varied significantly between those two periods *(p* = 0.035, i-test). In stress-trained rats, the fastest metabolic rates for Epi and NE were measured at 21.7 and 12.1 min, respectively *(**Figure 3C**).*

**Table 1.** The plasma miR-122 metabolic rates under different events

| Treatment | Half-life of Inducer<br>in circulation | plasma miR-122 Metabolic Rate |  |
| --- | --- | --- | --- |
|  |  | Average (*n*) | Shortest |
| Saline |  |  |  |
| 1st injection | n/a | 43.5 ± 14.5 (7) | 29.45 |
| 2nd injection | n/a | 18.1 ± 11.2 (3) | 11.2 |
| CDDP 9 mg/kg | >1800.0 <sup>a</sup> |  |  |
| 1st CDDP 0 - 4 hr |  | 41.3 ± 25.9 (3) | 19.8 |
| 1st CDDP 4 - 48 hr |  | 559.8 ± 266.7 (7)<sup>b</sup> | 120.8 |
| 2nd CDDP 0 - 48 hr |  | 608.9 ± 11.4 (2) | 600.9 |
| CDDP 30 mg/kg | >1800.0 <sup>a</sup> |  |  |
| 1st CDDP 0 - 4 hr |  | 278.0 ± 0.0 (1) | 278 |
| 1st CDDP 4 - 48 hr |  | 281.4 ± 96.3 (6) | 147.8 |
| 2nd CDDP 0 - 48 hr |  | 473.1 ± 397.1 (3) | 182 |
| Epinephrine | 3.5 <sup>a</sup> | 56.6 ± 2.1 (4) | 21.7 <sup>c</sup> |
| Norepinephrine | 2.0 <sup>a</sup> | 12.1 ± 0.0 (2) | 12.1 <sup>c</sup> |
| Dopamine HCl | 2.0 <sup>a</sup> | 96.6 ± 16.4 (3) | 82.4 <sup>c</sup> |
| Dexamethasone | 190.0 <sup>a</sup> | 1404.8 ± 1196.9 (6) | 418.1 <sup>c</sup> |

a: From the published clearance rate in venous plasma(Goldstein, *et al* 2003).
b: One outlier of 5174 min was excluded, if included, the average was 1136.6 min.
c: Tests on stress-trained rat models.
\*: Calculation based on the reduction of plasma miR-122 between two closest collection point.
Mean ± SD, unit: minute.

### The interday variations of human plasma miR-122 concentrations

To investigate interday variations, 12 volunteers composed with 6 healthy individuals, 2 Chronic Hepatitis B (CHB) carriers and 4 subjects with fatty livers (Table S2), were recruited in Sichuan Cancer Hospital. Five milliliters of venous blood were collected daily from each subjects between 9:00am to 12:00am for a 4-day period. As stress-relief measure, a 15-minute resting period was enforced before blood collection. The average interday levels of plasma miR-122 of each subject remained constantly with CV below 20.2% during the 4-day period *(**Figure 5A**).* Comparing each day value to 4-day mean of each individual, the interday variances of plasma miR-122 levels of healthy individual and individuals with chronic liver diseases were relatively invariant at daytime. More importantly, the relatively stable daytime plasma miR-122 levels were more dynamic in detection range comparing to serum ALT, AST, and Tbil.

**Figure 5.**
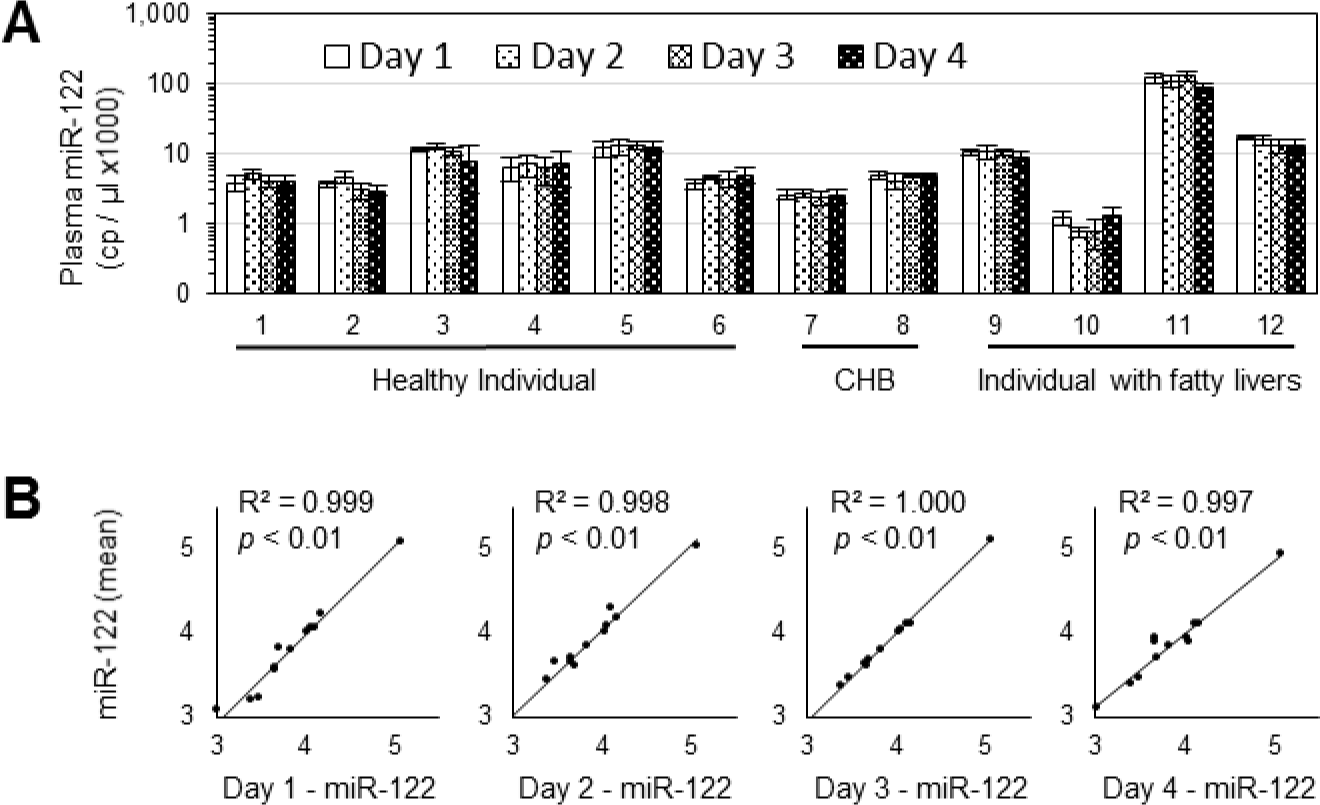
Interday variation of plasma miR-122 expression in 12 volunteers. (A) The daily plasma miR-122 concentrations at a 4-day period from each individual (Mean ± SD); (B) Pearson’s correlation analysis on the log-transformed plasma miR-122 concentrations of a given day vs log-transformed 4-day means of each individual. n = 4.

## DISCUSSION

Plasma contains three catecholamines: DA, Epi and NE. NE was identified as the main neurotransmitter in sympathetic nervous system regulating the cardiovascular system(Goldstein, *et al* 2003) while DA and Epi were related to acute stress responses. The plasma catecholamines were quickly converted to inactive forms of metabolites in liver through hepatic extraction by a set of enzymes(Eisenhofer, *et al* 2004). With the plasma-direct miRFLP assay, we showed that significant elevations of plasma miR-122 levels by catecholamine administration in stress-trained rats. Accordingly, acute stresses were identified as the major confounding factors to influence the plasma miR-122 measurements in naïve rats. Those data implicated that the degradation of physiological plasma catecholamines was linked to the basal levels of plasma miR-122.

The link between the basal plasma levels of ALT and circulating catecholamines has not been reported so far. The diurnal variation comparison revealed synchronized circadian rhythms of miR- 122(Ludwig, *et al* 2016), ALT(Cordoba, *et al* 1998), Epi and NE(Bondanelli, *et al* 1999, Linsell, *et al* 1985) in plasma samples. Plasma NE followed a diurnal rhythm of cosinor fit with relatively stable levels during daytime in healthy individuals(Bondanelli, *et al* 1999). Plasma Epi followed the same diurnal rhythm as plasma NE with much lower basal levels and significant fluctuations during daily activities due to emotional responses. Since hepatocellular miR-122 releases were promoted by circulating catecholamines and shared with the same diurnal rhythms, we hypothesized that the basal levels of plasma miR-122 and ALT might have reflected plasma catecholamine levels in healthy individuals, which could be easily influenced by stress and confounded by their half-lives. In comparison to the slightly decreasing plasma NE during daytime, plasma ALT levels increased up to 45% in afternoon than in morning in patients with cirrhosis(Cordoba, *et al* 1998). The morning measurements of serum ALT were closer to the plasma catecholamine basal levels than their afternoon values because of long ALT half-life. Our data indicated that the plasma miR-122 concentration were unaltered during daytime from 9:00am to 5:00pm in stress-trained rats *(**Figure 4A**).* In this sense, plasma miR-122 may be more dynamic than ALT in revelation of liver changes in “real-time” owning to their shorter half-lives and narrower intraday variances. The stress-free plasma miR-122 measurements may reflect the plasma catecholamine basal levels more closely for healthy rats and the severities of liver injuries.

The diurnal variance of plasma miR-122 levels was reported with up to 2-fold difference between daytime peak and nocturnal sleep(Heegaard, *et al* 2016). Comparison of plasma miR-122 diurnal variations between healthy individuals and patients, may reveal the mechanistic aspect of some liver diseases. For example, patient with cirrhosis is accompanied by higher plasma miR-122 which could be explained either as the leakages of miRNAs from hepatocytes or sensitized hepatocytes by cirrhosis. Useful clues could be raised by the comparison of diurnal concentrations of plasma miR-122 from cirrhosis patients.

The half-life of plasma miR-122 is still unclear in human but it was reportedly much shorter than that of serum ALT. We observed the fastest metabolic rates of ~12.0 min for plasma miR-122 in the second siHD phase of naïve rat and similar rates in stress-trained rats. This estimate is in agreement with a recent report with estimated plasma miR-155 half-life of 25 min(Bala, *et al* 2015). The plasma miR-122 existence in protein- or exosome-associated forms was also an important influence factor on their metabolic rates. Because of its fast metabolic rates, the predominant plasma miR-122 released by catecholamines might be in protein-associated form. The fast metabolic rate enables closer monitoring of liver injuries by plasma miR-122 than ALT. By substituting serum ALT with plasma miR-122, more accurate assessment of hepatocytes dying in an acute DILI event could be achieved as suggested in a recent mechanistic model(Howell, *et al* 2014).

Our data indicated that acute stress may have profound impact on miRNA release from hepatocytes at least. The correlated elevations of miR-92 with miR-122 in plasma samples under stresses revealed the potentially unselective nature of hepatocellular miRNA release. With hundreds of miRNAs expressed in hepatocytes, the stress management becomes a concern for any diagnostic application based on circulating miRNAs. For objective basal catecholamine measurement, a protocol was suggested in which the subject has been undisturbed in a quiet environment for 20 min before blood collection(Hart, *et al* 1989). This protocol was adopted in our interday variance study and proven to be effective in eliminating stress-induced miR-122 releases. This simple stress-relief measure was recommended as a prerequisite before blood collection of clinical applications.

## SUPPLEMENTAL INFORMATION

Supplemental Experimental Procedures, four figures and two tables can be found in Supplemental Information.

## ACKNOWLEDGMENTS

The authors thank Dr Rachel Church, Dr Paul Watkins and Dr Tao Lu for comments on the manuscript. Funding: This work was supported in part by Grants: 13ZZYB806GX-06 and 2014 Early R&D Project from the Science and Technology Bureau of Sichuan to Chengdu Nuoen Biotechnologies, Ltd. COMPETING INTEREST

J.Z., W.Z., Z.W., D.L. and K.X. are employees of Chengdu Nuoen Biotechnologies, Ltd.; all others declared none.

## AUTHOR CONTRIBUTIONS

All authors confirmed that they have contributed to the intellectual content of this paper with final approval of the publication. D.L., J.Z., J.H., B.X., H.Y. and K.X conceived the study and prepared the manuscript. X.P., Q.L., Z.Z., G.X., J.K., B.X., H.Y. and K.X. designed the experiments and analyzed data. W.Z., Z.W. and K.X. developed the miRFLP assay. X.P., J.Z. and W.Z. provided experimental ideas and technical support. X.P., X.H., X.S. and H.Y. performed animal experiments and data analysis. Q.L., S.L., S.Y., Z.Z., G.Z., L.L., T.Y. and J.Z. collected clinical samples and performed analyses on clinical data.

## Materials and Methods

### AILI mouse experiment

To investigate the correlation of AILI and plasma miR-122 concentrations, 68 female BALB/c mice (7-week-old, weighted 18.3 to 21.2 g) were acclimated for 7 days before experiments. The mice were randomly grouped (n = 4/group) and fasted overnight with free access to water. APAP (Sigma, Mouse LD50 = 338 mg/kg, oral) at 300 mg/kg (low-dose), 1,000 mg/kg (medium-dose), and 2,000 mg/kg (high-dose) were administered orally. Terminal bloods were collected at 2, 8, 24, 48, 72 hours (h) after APAP administration. Four mice were fed with placebos as negative control. Additionally, four mice were i.v. injected with 100 μl of saline 2-h prior to blood collection due to the original i.v. administration planning. The plasma miRNAs (miR-122, miR-192, miR-92 and miR-451) were quantitated simultaneously by a multiplexed plasma-direct miRFLP assay as reported previously(Xie, *et al* 2015, Zhu, *et al* 2017).

After blood collection, mice were sacrificed by cervical dislocation. Mouse livers were fixed in neutralized formalin and stained with hematoxylin and eosin (H&E). Histological analysis was performed by an experienced pathologist single-blindly as paid service by West China Medical College, University of Sichuan. and histopathological grades were scored according to the method reported previously(Park, *et al* 2016). The mice of high-dose groups were excluded from histological analysis for the narrow dose difference between medium- and high-dose APAP.

### Rat experiment with CDDP treatments

CDDP (Sigma, Rat LD50 = 33.8 mg/kg, intravenous) was used to verify that if siHD could be induced in naïve rats. Twelve female Sprague-Dawley (S/D) rats (7-week-old) were randomly separated into 4 groups and assigned letters from ‘A’ to ‘L’. The baseline samplings were taken at 0 and 24 h. At 48h, saline (or CDDP at 9, 19, and 30 mg/kg) was i.v. injected and blood samples were collected at 0, 20, 40, 60, 120, 240 and 320 minutes after saline or CDDP administration. At 96h, the routine of drug injection and blood samplings was repeated. All rats were lightly sedated with ether-inhalation for motion-restraining and 50 to 100 |il of blood were collected at each time point to minimize the impact on rat health. Plasma miR-122 concentrations were measured by the plasma-direct miR-122 miRFLP assay (***Figure 2 A to* l**). The technical specification of the assay was verified according to the Approved Guideline (EP5-A2) by Clinical And Laboratory Standards Institute (***Figure S4 A and* B**).

### Stress-trained rat model

The perception of stress was based on experience and could be reduced after repeated “Hits”(McEwen 1998). Based on the observation of animal adaptation to the repeat stress- stimulations in the negative control rats ‘A’, ‘B’ and ‘C’ *(**Figure 2 A to C**)*, a stress-trained rat model was developed to overcome the stress influence on plasma miR-122 measurement by the activities of injection and blood collections. Realizing that needle puncture during blood collection was also a source of stress, a routine, composed of a saline injection and four blood collections at 0, 2, 4, 8h after saline injection, was developed. Six female S/D rats (7-week-old) were trained with the routine every 2-3 days, and three routines a week (Monday, Wednesday and Friday), by a dedicated male operator. To minimize potential burdens on rat health, 50 to 100 μl of blood samples were collected each time. MiR-122 content in plasma sample from each time point was measured by the plasma-direct miR-122 miRFLP assay. After three routines, plasma miR-122 was determined and the model was established when there were no more plasma miR-122 elevations. From Day 8, saline was replaced by 100 μl of solutions containing different drugs, respectively. Blood samples were collected and analyzed according to the drug administration (***Figure 3 A to* C**).

### Stress stimulated liver injury in naïve rats

To investigate the roles of saline, ether-inhalation, injection and blood collection (needle puncture) in siHD, six naïve female S/D rats (7-week-old) were separated into three cages and treated subsequently with different routines including: a) saline injection with ether-inhalation; b) blood collection with ether-inhalation; c) saline injection without ether-inhalation. One hundred microliters of saline were injected to rats in group 1 and group 3. Fifty to one hundred microliters of blood were collected and analyzed before, and at 0.33, 0.66, 1, 2, 4 and 8 h after saline administration. No saline-injection was performed on rats in group 2 and blood collections were performed as other groups. The rats were handled in treatment orders exactly to examine the involvement of emotional stress.

### Plasma-direct miRFLP assay

We utilized modified plasma-direct miRFLP assay with noise-reduction adapters to quantitate the absolute concentration of miRNA from plasma samples directly. The detailed method was reported previously(Xie, *et al* 2015, Zhu, *et al* 2017). The absolute copy numbers of plasma miRNAs were calibrated against a serial dilution of miRXplore Universal Reference(Bissels, *et al* 2009).

Additional Experimental Procedures are included in Supplemental Information.

